# PDB ProtVista: A reusable and open-source sequence feature viewer

**DOI:** 10.1101/2022.07.22.500790

**Authors:** Mandar Deshpande, Mihaly Varadi, Typhaine Paysan-Lafosse, Sreenath Nair, Damiano Piovesan, Saqib Mir, Aleksandras Gutmanas, Silvio C. E. Tosatto, Sameer Velankar

## Abstract

The PDB ProtVista is a reusable and customisable sequence feature viewer that provides intuitive and detailed 2D visualisation of residue-level annotations while supporting interactive communication with 3D viewers. The Protein Data Bank in Europe (PDBe) team develops and maintains PDB ProtVista. Several public web services use it to display structural and functional annotations such as macromolecular interaction interfaces, intrinsic disorder predictions, sequence variants, and sequence conservation. The PDB ProtVista is freely available from https://github.com/PDBeurope/protvista-pdb. We provide extensive documentation and step-by-step user guides on integrating PDB ProtVista with existing web applications and 3D molecular viewers. We also offer examples of displaying the user’s custom data and functional annotations for PDB and UniProt entries powered by a rich set of PDBe API endpoints.

## Introduction

Identifying the biological functions of macromolecules is instrumental in understanding and potentially modulating cellular processes [1]. A widespread approach to unveiling these molecular mechanisms is to try and gain insights from the structural representations of proteins and nucleic acids [2,3]. While models can be helpful independently, macromolecular structures become more interpretable when combined with functional and biophysical annotations [4,5].

In 2018 we established the Protein Data Bank in Europe -Knowledge Base (PDBe-KB), which integrates the atomic coordinates of macromolecular structures in the Protein Data Bank (PDB) with annotations provided by data resources and research teams from across the globe [5]. While PDBe-KB offers a comprehensive resource of the biological context of protein structures, the complexity of these data makes it challenging to interpret the annotations and gain biological insights unless advanced and yet easy-to-use data visualisation tools are readily available to the scientific community [6]. A powerful and intuitive way of visualising annotations is to use two-dimensional (2D) sequence feature viewers to get an overview of all the annotated amino acid and nucleic acid residues for a particular macromolecular sequence [7]. There are several sequence feature viewers, some more general-purpose such as Feature-Viewer [8], and others more specialised, such as Mason [9] or pViz.js [10]. However, these sequence feature viewers were not optimised for the specific task of displaying structural annotations in 2D with seamless integration to 3D molecular viewers; a use case that is fundamental for data resources such as the PDB [4].

Here, we present PDB ProtVista, a 2D sequence feature viewer with native support for interactions with three-dimensional (3D) molecular graphics viewers. PDB ProtVista builds on and expands the Nightingale library [7], which is a collection of web components co-developed by our team, UniProt [11] and InterPro [12]. PDB ProtVista is open-source and available from https://github.com/PDBeurope/protvista-pdb.

Our expanded version of ProtVista is optimised for displaying annotations of protein and nucleic acid structure annotations. It includes out-of-the-box support for two-way interactions with the 3D viewer, Mol* [6]. Data resources such as PDBe (https://pdbe.org), PDBe-KB (https://pdbe-kb.org), CamKinet (http://camkinet.embl.de/v2/home/) and ProKinO (https://vulcan.cs.uga.edu/prokino) use PDB ProtVista to visualise annotations and map them interactively onto the 3D models [5,13,14].

## Materials and methods

PDB ProtVista extends and tailors the core ProtVista web components [7] to create an implementation specifically for displaying structural annotations found at PDBe and PDBe-KB. It is a native JavaScript (ES6) application which depends on the D3.js library to render its data visualisation. Its source code is openly available to the scientific community at GitHub: https://github.com/PDBeurope/protvista-pdb.

PDB ProtVista provides three data visualisation modes, called “tracks”, to support various types of sequence and structure annotations: i.) a “segment” track; ii.) a “variants” track, and iii.) a “sequence logo” track. The “segment” track, the most frequently used visualisation mode, displays block representations of either a single position in a sequence or a continuous range of residues. PDBe and PDBe-KB use this visualisation mode to display annotations such as structural domains [15,16], consensus disorder predictions from the MobiDB data resource [17] or continuous variables such as backbone flexibility propensity [18].

Amino acid variants can be displayed using a dedicated data visualisation track that supports displaying distinct sequence positions and their corresponding meta-information, such as the biological effects of mutations. PDBe and PDBe-KB use this track to display variants data from UniProt and sequence variants and their predicted structural and functional consequences provided by Missense3D [19] and FoldX [20].

Finally, PDB ProtVista can show sequence conservation as a histogram together with the probability that a particular amino acid appears at a given position using a sequence logo track. ProtVista shows the sequence logo when the user zooms in on the displayed sequence, and 150 or fewer residues are visible. We set this threshold to improve the loading performance and the user experience. Each amino acid is coloured using its electrochemical properties as defined by the Clustal X colour scheme [21]. PDB ProtVista sorts the amino acids according to their properties for each position by default. Users may also sort the sequence logo by probability using the corresponding option on the left-hand side of the track.

The PDBe weekly release process includes the sequence conservation calculations that power this data visualisation track, ensuring that up-to-date data are displayed. The weekly process generates sequence conservation information for every PDB entry and corresponding UniProt accessions when sufficient sequence data is available in UniProt. The protein sequences are aligned against the UniProt database to obtain multiple sequence alignments, and the process generates Hidden-Markov Model profiles using the HMMER suite [22]. The data pipeline then calculates amino acid probabilities and conservation scores from the profiles obtained. Users can download the multiple sequence alignments (MSAs) corresponding to the displayed sequence logo by clicking on the “Download MSA” button in PDB ProtVista.

There are two main modes of integrating PDB ProtVista into existing services. Researchers and software developers can display their custom annotations or use application programming interface (API) endpoints that serve PDB and PDBe-KB data in ProtVista-compliant JSON (JavaScript Object Notation) format. We provide interactive demos on displaying data through both data access mechanisms, comprehensive documentation, user guides and linking PDB ProtVista to the 3D-viewer Mol* at https://github.com/PDBeurope/protvista-pdb/wiki.

## Results

PDBe-KB integrates macromolecular structure data with functional and biophysical annotations in a comprehensive knowledge graph [5]. The data in this archive is complex and diverse, necessitating advanced and easy-to-use data visualisations to help the scientific community gain valuable insights regarding the biological role of macromolecules. We created PDB ProtVista, a 2D sequence feature viewer consisting of reusable, framework-agnostic web components to provide an overview of structural and sequence annotations with a focus on supporting seamless communication with the 3D-molecular graphics viewers such as Mol^*^ [6].

We demonstrate the functionality of PDB ProtVista through the example of Calpastatin (UniProt accession P27321), an inhibitor of the calcium-dependent, non-lysosomal cysteine protease Calpain (Figure 1).

**Figure 1.**
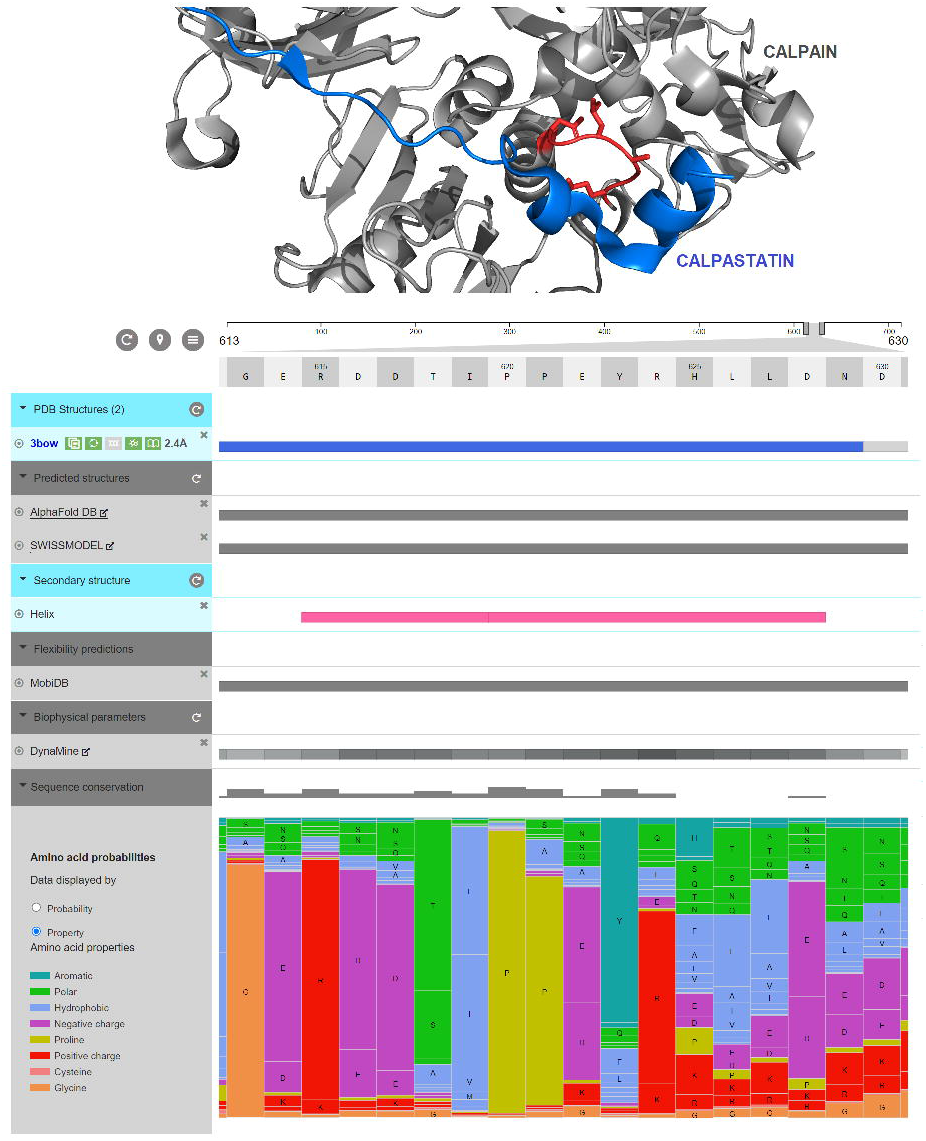
-PDB ProtVista is displaying annotations for Calpastatin. Calpastatin is the native inhibitor of the protease Calpain. Using publicly available data, PDB ProtVista can display all the related PDB entries and predicted models from AlphaFold DB and SWISS-MODEL. It can also show the observed secondary structure elements (where the pink rectangle represents a helix) and all the amino acid positions that interact with Calpain (blue rectangles). PDB ProtVista can also show computationally derived annotations (in grey), such as consensus disorder predictions from MobiDB, backbone flexibility propensities by DynaMine and sequence conservation propensities complemented by a sequence logo. Examining these annotations highlights the region where Calpastatin adopts a helical structure. This region bends out over the catalytic site of Calpain (coloured red in the figure above) to avoid enzymatic cleavage. These PDB ProtVista visualisation tracks are used at https://www.ebi.ac.uk/pdbe/pdbe-kb/proteins/P27321 to display Calpastatin annotations.

Using publicly available API endpoints [23], PDB ProtVista can list all the known structures of Calpastatin in the PDB and predicted models from AlphaFold DB and SWISS-MODEL [24,25]. It also displays information on the observed secondary structure elements, complemented by consensus disorder predictions from MobiDB and backbone flexibility predictions from DynaMine. PDB ProtVista can also highlight the interaction interfaces between Calpastatin and Calpain and display sequence conservation and sequence logo data. Together, these data visualisation tracks show that Calpastatin is an intrinsically disordered protein, according to MobiDB, and it has regions that may adopt context-dependent secondary structure, according to DynaMine. These regions adopt short helices in the Calpain/Calpastatin complex (PDB structure: 3bow)[26]. PDB ProtVista also highlights that the sequence conservation propensity is higher for residues that do not directly interact with Calpain. This seemingly counterintuitive observation reflects the actual mechanism by which Calpastatin inhibits Calpain. The inhibitor protein binds to a long cleft on the catalytic subunit of Calpain. To avoid being cleaved by Calpain, Calpastatin loops out by forming a short helical secondary structure element, explaining why the sequence conservation of this region is higher than that of the flanking macromolecular interaction interfaces.

In addition to providing such functional insights through the 2D overview of annotations, an essential feature of PDB ProtVista is its capacity for seamless interaction with the 3D molecular graphics viewer, Mol*. For example, users can compare intrinsic disorder predictions with the observed interaction interfaces and secondary structure elements in 2D and 3D. The MobiDB data resource provides these up-to-date consensus disorder annotations that PDBe-KB integrates weekly with the core PDB data [5]. Two-way communication between Mol* and PDB ProtVista makes it easy to interactively identify regions that are likely to adapt semi-stable secondary structures when bound to their interaction partners, such as Calpastatin’s case described above.

## Discussion

Macromolecular structures are most useful in understanding the biological function of a molecule when used together with functional and biophysical annotations. These annotations can provide the biological context of proteins and nucleic acids, from ligand binding sites to the effects of mutations and from backbone flexibility predictions to macromolecular interaction interfaces [17–19].

Displaying a rich set of annotations directly on the 3D representation of a macromolecule is a complex problem, and such data visualisations may become difficult to interpret. A popular approach is to display annotations in a linear, 2D way. These data visualisation tools are collectively referred to as sequence feature viewers, and they make data more accessible and more interpretable [7–10]. These viewers traditionally focus on and excel in displaying sequence features and sequence-derived annotations, but they are often less optimal for displaying structure-based annotations, and in particular, they lack support for interacting with 3D molecular viewers.

Here, we presented PDB ProtVista, an expanded, specialised version of the collaboratively developed ProtVista viewer. We created this data visualisation application with the specific goal of viewing structural and functional annotations in 2D while being able to concurrently map the annotations onto 3D visualisations. Our data resources, PDBe and PDBe-KB, rely on this sequence feature viewer. Other data providers, such as CamKinet, have also adopted this open-source tool to display annotations for protein and nucleic acid structures.

With more protein structures available to the scientific community than ever before [24], it is timely to have a sequence feature viewer dedicated to the annotations of these macromolecule models in a way that can help researchers gain novel insights regarding the functions and biological role of proteins.

## Acknowledgements

We thank the UniProt and InterPro teams for the collaborative development of the core ProtVista sequence feature viewer. We are also indebted to the Velankar team, who helped test and improve PDB ProtVista. We would like to acknowledge funding from ELIXIR [IDP implementation study], the Biotechnology and Biological Sciences Research Council via the 3D-Gateway [BB/T01959X/1], Genome3D (BB/N019172/1), and FunPDBe [BB/P024351/1] grants, the Wellcome Trust [104948] and the European Molecular Biology Laboratory-European Bioinformatics Institute who supported this work.

